# Evolution of the genetic architecture of local adaptations under genetic rescue is determined by mutational load and polygenicity

**DOI:** 10.1101/2020.11.09.374413

**Authors:** Yulin Zhang, Aaron J. Stern, Rasmus Nielsen

## Abstract

Inbred populations often suffer from heightened mutational load and decreased fitness due to lower efficiency of purifying selection at small effective population size. Genetic rescue (GR) is a tool that is studied and deployed with the aim of increasing fitness of such inbred populations. The success of GR is known to depend on certain factors that may vary between different populations, such as their demographic history and distribution of dominance effects of mutations. While we understand the effects of these factors on the evolution of overall ancestry in the inbred population after GR, it is less clear what the effect is on local adaptations and their genetic architecture. To this end, we conduct a population genetic simulation study evaluating the effect of several different factors on the efficacy of GR including trait complexity (Mendelian vs. polygenic), dominance effects, and demographic history. We find that the effect on local adaptations depends highly on the mutational load at the time of GR, which is shaped dynamically by interactions between demographic history and dominance effects of deleterious variation. While local adaptations are generally restored post-GR in the long run, in the short term they are often compromised in the process of purging deleterious variation. We also show that while local adaptations are almost always fully restored, the degree to which ancestral genetic variation comprising the trait is replaced by donor variation can vary drastically, and is especially high for complex traits. Our results provide considerations for practical GR and its effects on trait evolution.

## 1. Introduction

Genetic rescue (GR) is a strategy used in conservation biology to increase fitness of an endangered inbred (recipient) population by introducing genetic variation from another (donor) population. GR is accomplished by assisted migration of individuals from closely related, healthy populations to the inbred imperiled population. This process naturally causes the replacement of local genetic variation in the recipient population with that of the donor population. Typically, only a small number of individuals are introduced in order to conserve local genetic variation (Whiteley, Fitzpatrick, Funk & Tallmon, 2015).

The strategy has now been practiced on many highly inbred populations from different taxa, including the Florida panther (Johnson et al., 2010), robins (Heber et al., 2012), guppies (Fitzpatrick et al., 2016), wood rats (Smyser et al., 2013), and adders (Madsen et al., 1999). In several cases, GR efficiently increased the absolute fitness of the inbred population and reduced inbreeding depression (Frankham, 2015). A famous example is the introduction of mountain lions from Texas of the sub-species *P. c. stanleyana* to the Florida *P. c. coryi* population, for which the number of *P. c. coryi* increased three-fold after only five years, with increased survival rates and a doubling of the heterozygosity (Johnson et al., 2010). A meta-analysis provided evidence that the beneficial effect of GR can persist through the F3 generation (Frankham, 2016). These empirical tests suggest that GR is a powerful conservation tool for increasing fitness in endangered inbred populations.

However, despite its promise, there is skepticism and caution towards the application of GR due to concerns about outbreeding depression and genetic homogenization (Bell, *et al*. 2019). In the case of the Florida panther, an estimated genetic replacement of 41% has been reported. In another case of the Isle Royale wolf, the immigration of one single male to Isle Royale caused a genetic replacement of 56% to the local inbred population within two generations (Adams et al. 2011). Hwang et al (2012) also reported a negative fitness effect after practicing GR with two species that are genetically highly divergent due to outbreeding depression.

Several different theoretical studies have been conducted to examine the expected efficacy of GR (Hedrick, Hellsten & Grattapaglia, 2016; Harris, Zhang & Nielsen, 2019; Tallmon et al., 2004; Frankham, et al., 2011). The dynamics of GR is complex, depending on, among other factors, the amount of gene-flow, the demographic model (e.g. effective population size), and the dominance coefficients of mutations. Harris et al (2019) showed that with a higher amount of introgression, the relative fitness of the recipient population recovers more quickly; however, this occurs at the cost of replacing an increasing proportion of the recipient’s ancestral genomes with those of the donor population. Demographic history of the recipient and donor populations also determines the dynamics of GR. For example, small effective population size (*N*_*e*_) limits the efficacy of natural selection; thus, in most cases admixture from a population with large *N*_*e*_ helps restore fitness (Harris et al., 2019). However, several studies have shown that demography and dominance of deleterious mutations have key interaction effects on the GR process (Harris et al., 2019; Kyriazis, Wayne & Lohmueller, 2019). For example, Kyriazis et al (2019) showed that fitness in populations with historically low *N*_*e*_ can be more robust to severe bottlenecks than those with historically large *N*_*e*_, as these populations are less efficient at purging recessive deleterious mutations.

Previous studies suggest that the genetic replacement caused by GR can be controlled if the amount of admixture is limited (Harris et al., 2019; Whiteley et al., 2015; Bell et al., 2019). However, whether local adaptation plays a role in GR remains an open question. Recently, Osmond & Coop (2019) investigated the population genetic signatures of selective sweeps under evolutionary rescue, i.e. the adaptive response and recovery from reduced absolute fitness due to environmental change. Also, Tomasini & Peischl (2019) investigated the effect of local adaptations on evolutionary rescue. But whether GR would lead to the loss of unique local adaptations, or whether local adaptations could affect the process of fitness restoration by GR, remain largely unexplored.

Here, we explore how the addition of linked locally-adaptive variation affects the GR process. Specifically, we explore the dynamics under GR of (1) a Mendelian trait, and (2) a polygenic trait under stabilizing selection with a shift in the optimum. Our results illustrate how the genetic architecture of adaptive traits evolve under GR, and how the dynamics of GR depends on the joint effects of demographic models and genetic factors such as dominance.

## 2. Materials and Methods

We simulate under two demographic models from Harris, et al. (2019), as illustrated in Figure 1. The simulations are conducted using SLiM (Haller & Messer, 2019). Model 1 (Fig. 1A) represents a population that undergoes a long-term bottleneck of 0.1 times the ancestral population size (*N*_*e*_=10^4^), which last for 16,000 generations. This demographic history represents a population that is inbred for a long period of time and is similarly to that estimated for Neanderthals by Prufer *et al*. (2014). Neanderthals represent a good example of a long-term inbred population where genomic analyses have discovered a substantial accumulation of deleterious alleles (Prufer *et al*., 2014). Model 2 (Fig. 1B) represents instead a population with an extreme, short-lived bottleneck with *N*_*e*_=10 that lasts for 20 generations, which might be more representative of many currently endangered species. Prior to the time of divergence, we conduct a burn-in phase of 44,000 generations. We simulate two modes of adaptation: a Mendelian trait, with only one adaptive site contributing to the trait, and a polygenic trait controlled by a large mutational target.

**Figure 1.**
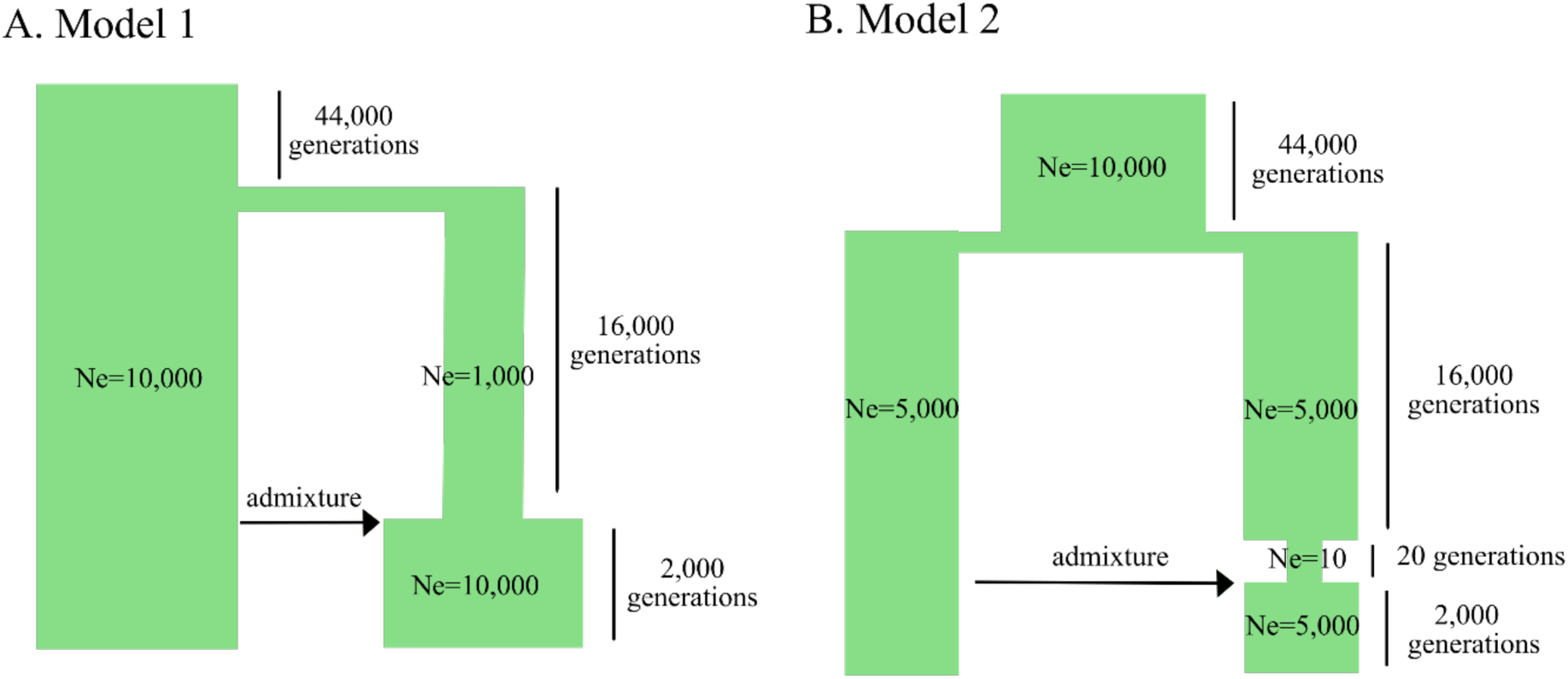
Two demographic models used for simulations. Time runs from top to bottom. Admixture happens at the generation right after the population size increases and lasts for only one generation. Population size changes are assumed to be discrete as depicted in the figure. Samples are taken from the inbred population on the right (p2) after the admixture up to 2,000 generations.

We simulate the Mendelian trait under two selection models: (1) a hard sweep, in which a rare additive beneficial mutation occurs after the split of the population, and (2) a soft sweep from a standing variant, in which an allele segregating neutrally leading up to the split is picked at random, and its selection coefficient is then changed so that the allele is then beneficial. Selection acts on the trait only after the split of two populations. For both models, we have examined different selection coefficients, *s* = 10^−4^, 10^−3^, and 10^−2^ for the adaptive mutation.

We simulate the polygenic trait under a model of stabilizing selection with Gaussian fitness. To model the effects of local adaptation in the recipient population we allow the phenotypic optimum in this deme to increase by some amount, immediately following the divergence from the ancestral population. With *V*_S_ as the variance of the fitness function (not to be confused with the variance in fitness among individuals), we simulated scenarios where the inbred population’s phenotypic optimum shifts by *δ* = 1, 2, 5 immediately after the split, while the phenotypic optimum remains 0 for the outbred population. We considered different selection strengths by setting the variance of the fitness function to be *V*_*S*_= 3,000 and 10,000. We assume genetic effects among loci are purely additive. Under this model, at equilibrium (phenotypic mean equal to the optimum), alleles are under under-dominant selection with *s* = *a*^2^/*V*_S_, where *a* is the effect of the allele, on the same scale on which the fitness function is defined (Simons, *et al*. 2018). In the transient phase after a large shift in the optimum, selection is approximately additive with *s* = *aδ*/*V*_*S*_ (Hayward & Sella, 2019). In order to ensure selection coefficients of causal SNPs are roughly *s* ∼ 10^−4^ − 10^−3^, in line with current estimates that SNPs ascertained for complex traits in humans have been under weak selection (Simons, et al. 2018), we draw the effects of causal alleles (*a*) from a standard normal distribution (i.e. mean 0 and variance 1). See Table S1 for more details.

In addition to the adaptive mutations described above, we also allow for accumulation of deleterious mutations assumed to be (1) additive (*h*=0.5), (2) partially recessive (*h*=0.1) and (3) recessive (*h*=0), where *h* is the dominance coefficient. To specify a set of simulation parameters realistic for mammals, we chose parameters estimated in humans for recombination rates and distribution of fitness effects. We use the UCSC exon map from the HG19 genome and the Distribution of Fitness Effect (DFE) on non-synonymous mutations estimated by (Eyre-Walker, Woolfit, & Phelps, 2006), assuming a non-synonymous mutation rate of 7×10^−9^ per bp/generation and log additive interactions among selected loci. A summary of the simulations is provided in Supplementary Table S1.

For all simulations, we have recorded fitness in the inbred population relative to that of the outbred population, the ancestry proportion in the inbred recovering population, the varying allele frequency of the adaptive mutation in the Mendelian model, and the fluctuation of mean phenotype in the stabilizing selection model.

## 3. Results

### 3.1 Selection on Mendelian traits

We simulated a Mendelian trait that is fixed for the derived (locally adaptive) allele in the recipient population, and fixed for the ancestral allele in the donor population. We varied the selection coefficient on the trait, the dominance coefficient of the linked deleterious variation, the admixture proportion during GR, and the demographic model (Model 1 vs. Model 2, see Fig. 1).

We investigated the effect of GR on fitness in the recipient population (i.e. hybrid fitness, Fig. 2 A-C) as well as on ancestral genome proportion (Fig. 2 D-F). Fitness is measured by taking the average of the fitness of offspring in the recipient population, and normalizing by the same quantity for the donor population. The fitness calculated in generation *T* is the fitness of parents (rather than offspring produced) in generation *T*. We found that, depending on the demographic model and dominance of deleterious variation, GR has drastically different success in terms of achieving rapid increase in hybrid fitness. For example, when deleterious mutations are partially recessive, GR is successful but somewhat slow for Model 2 (Fig. 2E). In Model 1, under the same scenario, fitness is not fully recovered even after 1000 generations post admixture with 10% admixture from the donor population (Fig 2B). By contrast, under a fully recessive load, fitness is restored extremely quickly under Model 2, provided sufficient admixture (1%), whereas with the same level of admixture in Model 1, fitness is not restored even in the long run (>1000 generations post admixture) (Fig 2C).

**Figure 2.**
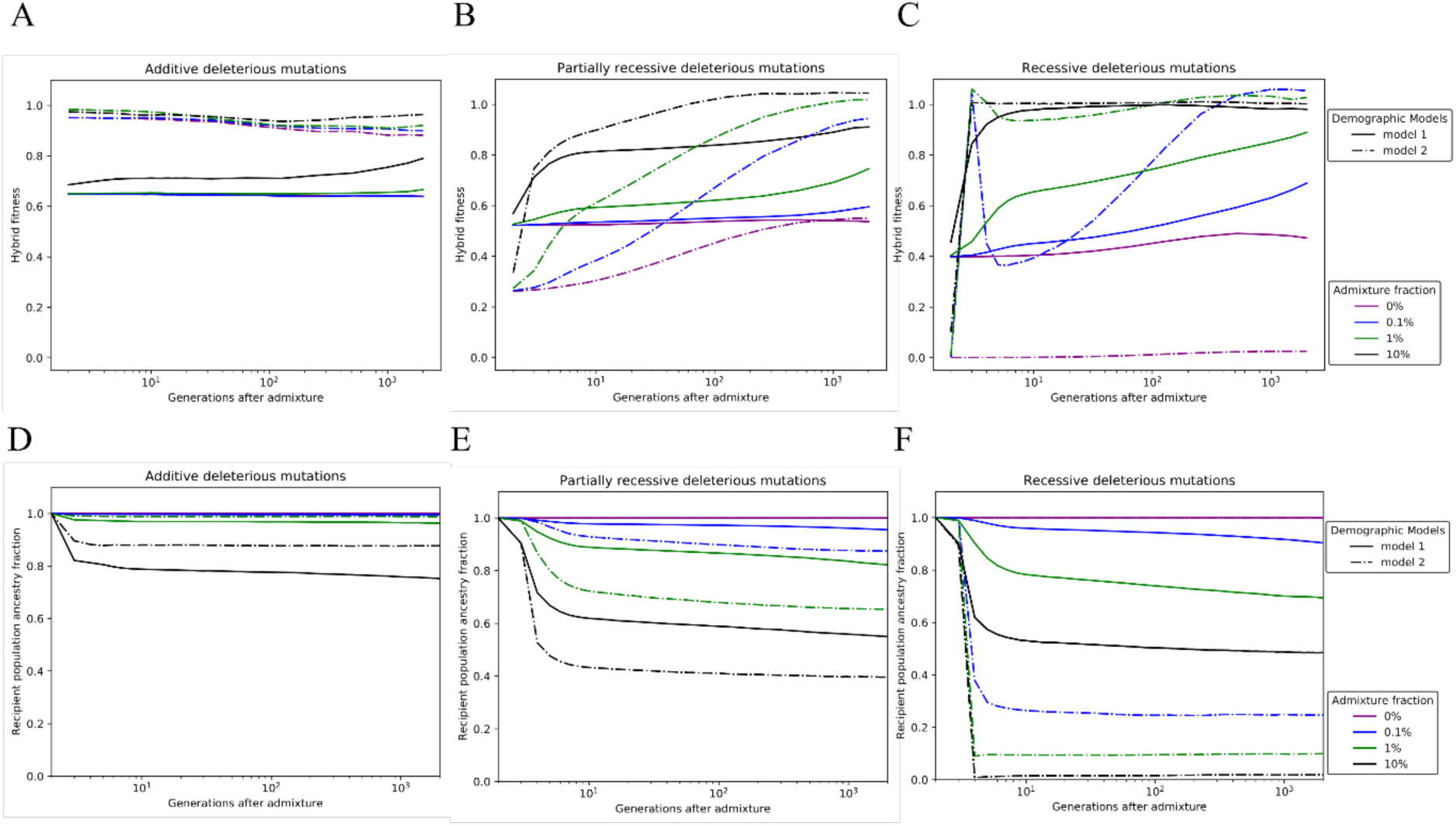
(A-C) Hybrid fitness change of the inbred population after admixture and (D-F) Recipient population ancestral genome fraction changes after GR with a Mendelian adaptive trait, under hard sweep selection model and demographic Model 1 (solid lines) and demographic Model 2 (dashed lines). The adaptive mutation is additive (dominance coefficient *h*=0.5) while selection coefficient is set differently (shown with different line styles in A-F). Deleterious mutations are assumed additive (*h*=0.5) in A, D, partially recessive (*h*=0.1) in B,E and recessive (*h*=0) in C,F.

Generally, we find that the lower the recipient fitness before admixture, the higher the amount of genomic replacement by the donor population in the long run (Fig. 2D-F). Furthermore, the more successful the GR is at restoring fitness, the higher the amount of genomic replacement in the long run (Fig. 2C,F). These conclusions are similar to those previously observed by Harris *et al*. (2019).

We also considered how ancestry and fitness evolve jointly in the recipient population (Fig. 3). Here, we show dynamics for a population under Model 2 with recessive deleterious variation. We found that in the first generation after GR, native ancestry is either 0 or 100% in the parents, where native ancestry is associated with much lower fitness (Fig. 3A). After one generation of admixture, a large proportion of offspring were inbred-outbred crosses, despite low admixture proportion (1%). Due to the large fitness advantage associated with outbred ancestry, these crossed individuals enjoyed much higher fitness than not only the inbred individuals, but also the non-crossed outbred individuals, because it is extremely rare for these crosses to be homozygous for recessive deleterious variation (it would require a recessive deleterious variant to segregate in both inbred and outbred populations since divergence up to the admixture). In the following generations, as ancestry proportions range between ∼30-90%, there is a clear trend of lower native ancestry incurring increased hybrid fitness (Fig. 3C,D)

**Figure 3.**
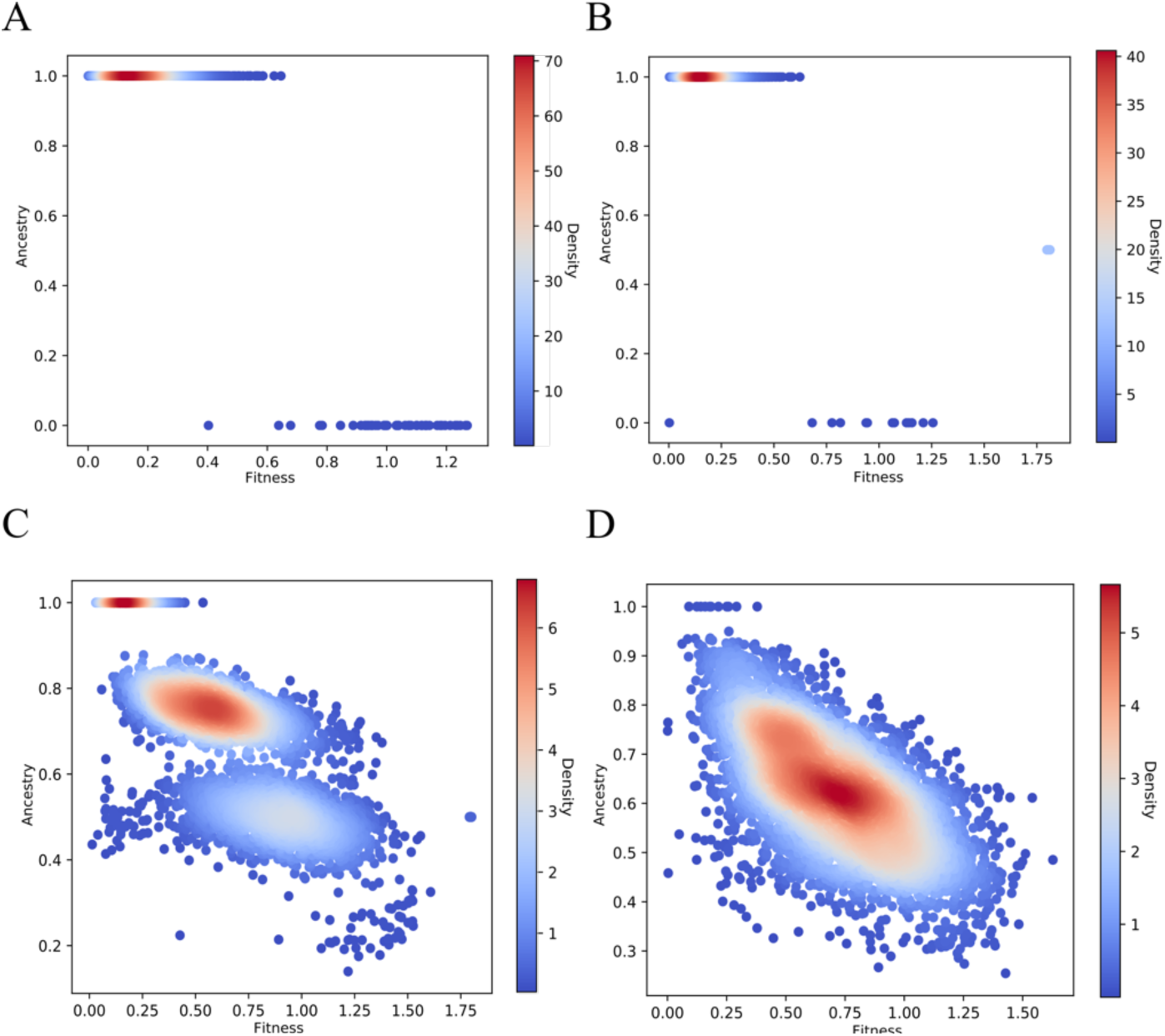
Relation between individual fitness and its ancestral genome proportion under demographic Model 2, with recessive deleterious mutations and admixture fraction of 1%. Each dot represents an individual, depicting relation between ancestry proportion and relative fitness (to the mean fitness of the outbred population) of each individual in the inbred recipient population. Figure A shows the population before mating with outbred individuals. Figure B indicates the first generation after admixture (e.g. F1), while figure C represents the second generation (e.g. F2) and figure D shows the third generation (e.g. F3).

A short term bottleneck (Model 2) does not increase or decrease the average number of mutations an individual carries. However, it will allow recessive deleterious mutations of strong effect, which were already segregating in the population, to increase in frequency and potentially go to fixation (while others are lost). Models of recessive mutations allow for much more standing variation of deleterious mutations, that potentially can increase in frequency during the bottleneck, than models of additive mutations (e.g. Fig. 2A vs 2C). GR is particularly effective in this case because the recipient population may have fixed strongly deleterious recessive mutations that can be purged immediately after GR (Fig. 2C). In the case of a constant low population size (Model 1), deleterious mutations (both in the recessive and additive model) will accumulate and can slowly go to fixation if they have weak effects, but one is unlikely to observe the same kind of strong effect of strongly deleterious recessive mutations going to fixation as you would in models with recessive mutations and a bottleneck (Fig. 2B-C).

Next, we looked at the dynamics of local adaptations under a Mendelian trait model (i.e., a single adaptive allele). We found that, in the short term (0-100 generations post-GR) the adaptive allele decreases in frequency following the loss of ancestral DNA fractions (Fig. 4; Fig. S1), which is the consequence of a selection-induced reduction in native ancestry after the admixture (Fig. 2D-F). However, while the ancestral genome proportion continued to decline slowly after 10 generations after GR (Fig. 2D-F), the adaptive allele generally increased in frequency after 10-100 generations when enough recombination occurred to break up linkage between the adaptive allele and the deleterious alleles. However, even for very strong selection (*s* = 0.01), it took hundreds of generations for the adaptive allele to reach high frequency in the population. In some extreme cases, with sufficiently high levels of admixture (10%), GR under Model 2 actually caused the adaptive allele to be lost with high probability after a total genetic replacement when the selection coefficient is sufficiently small (≤0.001) (Fig. 4F).

**Figure 4.**
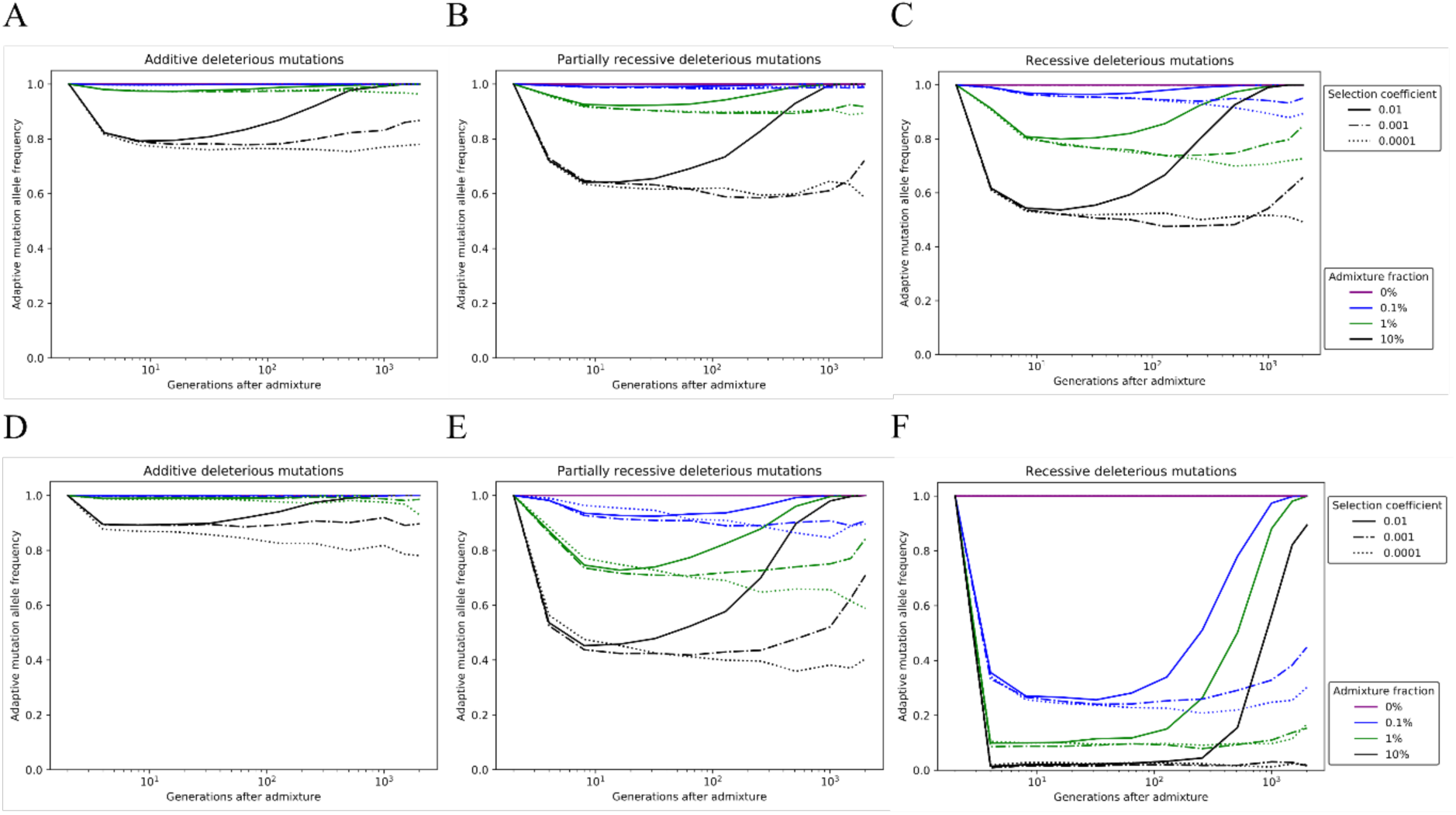
Allele frequency changes after admixture for a Mendelian trait under hard sweep selection model and (A-C) demographic Model 1 (D-F) demographic Model 2. The adaptive mutation is additive (dominance coefficient *h*=0.5) while selection coefficient is set differently (shown with different line styles). Deleterious mutations are assumed additive (*h*=0.5) in A, D, partially recessive (*h*=0.1) in figure B,E and recessive (*h*=0) in C,F.

We also examined the joint effects of dominance coefficients of linked deleterious variation and demographic history on the efficacy of GR. Under both model I and II, we saw a greater degree of genetic replacement (Figure 2D-F), leading to a greater reduction in the frequency of the adaptive allele, as deleterious mutations become more recessive (Fig. 3 & S1). For example, there was a smaller short-term reduction in allele frequency of the adaptive allele under Model 2 (Fig. 4D-F, Fig. S1 D-F, relative to Model 1 [Fig. 4A-C, Fig. S1 A-C]) but a larger reduction for partially recessive/recessive deleterious variants (Fig. 4B,C,E,F, Fig. S1 B,C,E,F), following the same pattern of the ancestral DNA proportion decline in those scenarios (Fig 4. G-H). However, the locally adaptive locus itself had only a minor effect on the recovery of relative fitness and reduction of ancestral DNA proportion of the recipient population (Fig. 4, 5, S1, S2, S3). We considered that sweeps from standing variation may have different patterns of linked deleterious variation around the adaptive allele; however, when simulating under alternative selection models we found no difference between the hard *vs* soft sweep models (see Fig. 4, S2, S3 vs Fig. S4, S5, S6).

**Figure 5.**
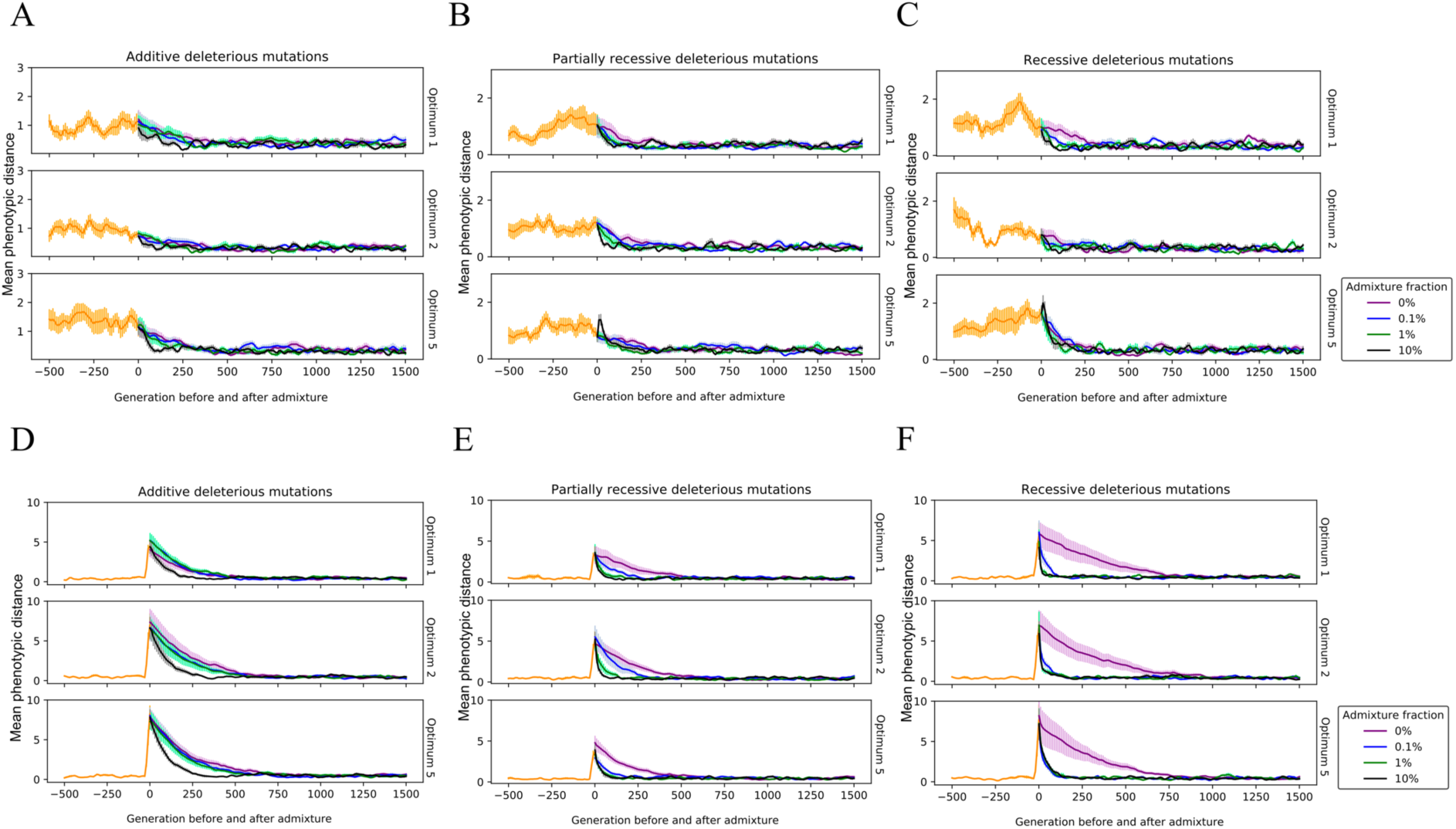
Mean phenotype distance from optimum over time. (A-C) Simulations under Model 1, (D-F) simulations under Model 2. Shaded bars signify 95% confidence intervals for the mean phenotypic distance.

### 3.2 Polygenic adaptation

We also simulated a polygenic trait under a model of stabilizing selection with a shift in the optimum (Fig. 5-7). Here we varied the strength of stabilizing selection on the trait (controlled by 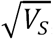, the ‘width’ of the fitness function), the size of the shift in the local optimum after population 2 diverges from population 1, *δ* = 0,1,2,5, measured in units of 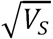), dominance coefficient of deleterious mutations, admixture fraction during GR as well as demographic model.

We examined two features of polygenic trait evolution: first, we evaluated the effect of GR on the perturbation of the adaptive phenotype from its optimum (Fig. 5, Fig. S7); we measure this by looking at the average distance of the population mean phenotype from the optimum. We also considered the extent of replacement of ancestral variation causal for the trait (Fig. 6); we measure this replacement by examining the relative proportion of genetic variance of the trait due to ancestral variation *vs* donor variation introduced by GR.

**Figure 6.**
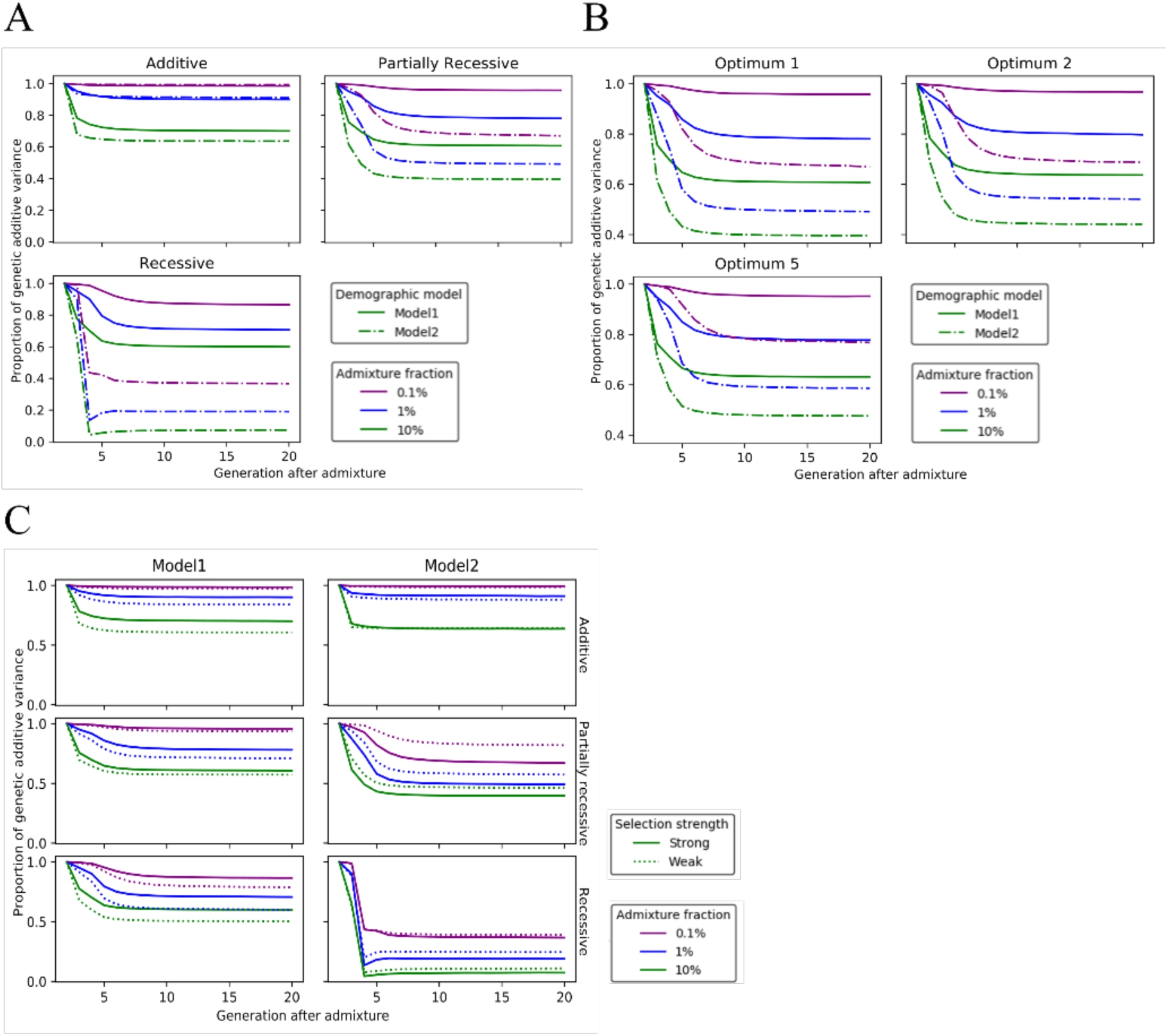
Proportion of genetic additive variance of a polygenic adaptation, recording mutations that originate from the recipient population for 20 generations after GR. (A) Scenarios under strong selection, with optimum of 1 for the recipient population and different dominance coefficients for deleterious mutations. (B) Scenarios under strong selection, with partially recessive deleterious mutations and different optimum for the recipient population;(C) Scenarios with optimum equals 1 and different selection strength for the adaptive trait. Each line represents the proportion of genetic additive variance contributed by variants, which compose the trait, from the ancestral genome of inbred population. Here, genetic additive variance is calculated as 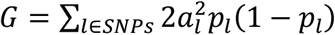, where *a*_*l*_ represents the effect of SNP *l*, and *p*_*l*_ its frequency.

We found that under Model 1, polygenic adaptations are not significantly affected by GR, as the trait’s evolution appears to follow the same trajectory regardless of admixture proportions or dominance coefficients of the deleterious load (Fig. 5A-C). However, under Model 2, we found that following rapid phenotypic drift from the optimum due to a severe bottleneck (Fig. S7), polygenic adaptations subsequently follow dramatically different trajectories depending on several factors (Fig. 5D-F): for example, GR allows the polygenic adaptation to recover to its optimal value much more quickly than without GR (Fig. 5F); and this effect is most pronounced under scenarios where there is fully recessive load (Fig. 5F), although it is still significant under a partially recessive load (Fig. 5E).

We also explored how the genetic basis of the polygenic adaptation in the recipient population is replaced by donor variation (Fig. 6). We quantify this using the proportion of the genetic variance attributable to standing variation in the recipient population just before admixture; genetic variance post-admixture is the sum of this quantity, plus genetic variance attributable to standing variation in the donor population just before admixture, plus that of *de novo* mutations occurring in the recipient population post-admixture (although this has negligible contributions over short timescales). Generally, we find that the genetic basis is quickly replaced due to GR, with >90% of the genetic variance being replaced with donor variation when GR is most successful; for example, under Model 2, especially when the deleterious load is recessive and the admixture fraction is high (Fig. 6A). Broadly, patterns of genetic variance replacement are consistent with patterns of ancestry replacement (Fig. 6 *vs*. Fig. 2D-F), with stronger replacement in situations where GR is more successful at recovering fitness. However, details of the local adaptation do affect the dynamics of how the genetic variance evolves; for example, when the optimal phenotype is more highly diverged in the recipient *vs* donor population, the fraction of the genetic variance replaced by the donor population is lower (Fig. 6B), because in this case donor individuals are more poorly adapted to the environment of the recipient population, and thus GR is countervailed by this force.

Lastly, we directly compared hybrid fitness trajectories under the Mendelian *vs*. polygenic trait models (Fig. 7). We found that, under the simulations models we considered, varying parameters controlling the local adaptation does not have any appreciable effect on hybrid fitness as an outcome of GR. In Fig. 7 we compare simulations of a Mendelian trait under strong selection *vs*. a polygenic trait under strong stabilizing selection (see Methods). The results are comparable both under Model 1 and Model 2 (Fig. 7A,B *vs* 7C,D) and under various levels of dominance (Fig. 7).

**Figure 7.**
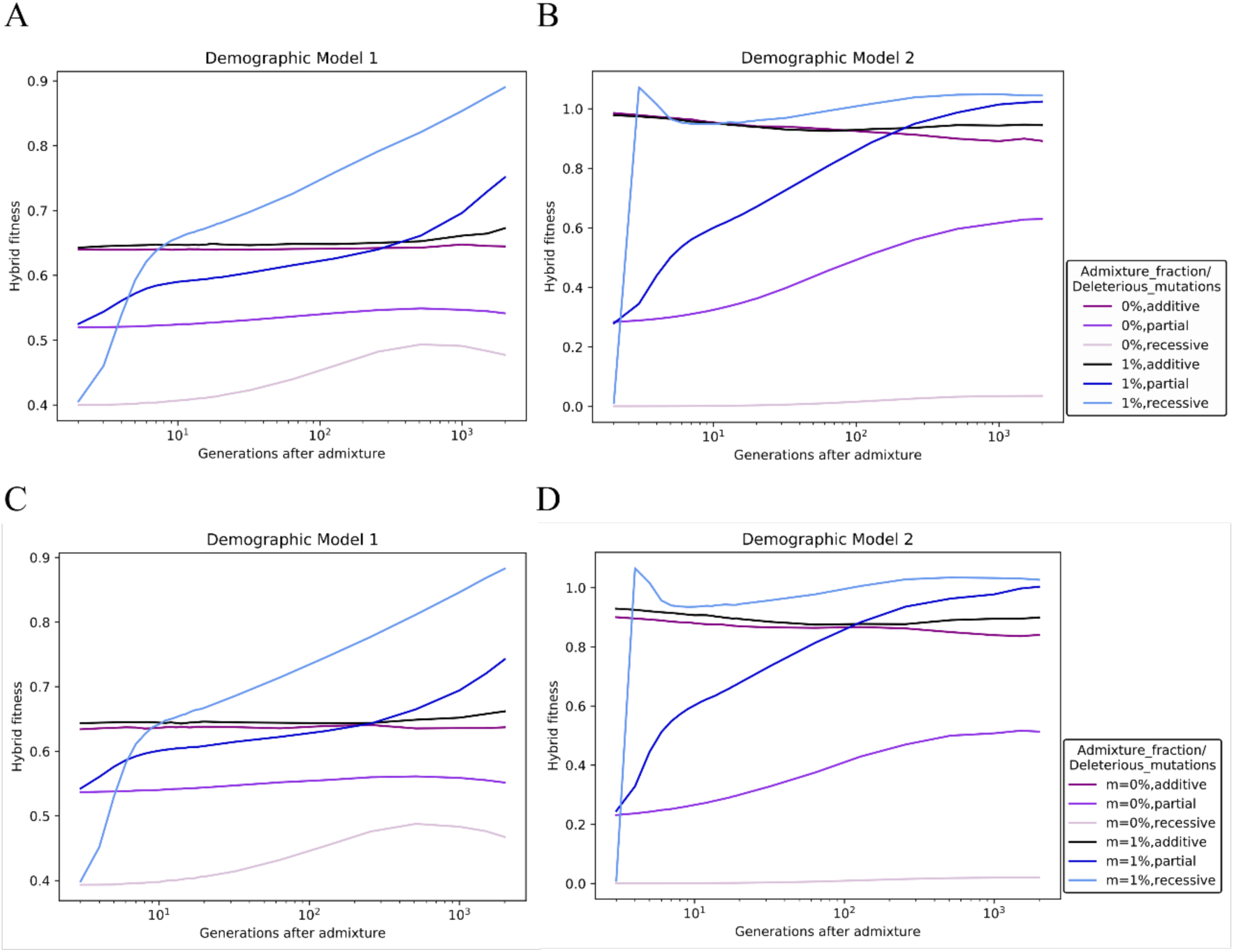
Hybrid fitness changes of the inbred population after admixture with (A-B) a Mendelian adaptive trait, under hard sweep selection model and (C-D) a polygenic trait under stabilizing selection with strong selection. Figure A and C are results of demographic Model 1 while B and D are that of demographic Model 2. Lines of different colors are indicating scenarios with different admixture fraction and dominance (of deleterious mutations).

## 4. Discussion

We have presented a population genetic simulation study that elucidates the dynamics of local adaptation and genetic rescue (GR). We considered various models of the selection strength and architecture of the adaptive trait, dominance of the mutational load, demography, and admixture. The results of our simulations show that when a locally adaptive trait consists of a single locus (e.g. a Mendelian trait), GR decreases the allele frequency in the short term. Dominance of the linked deleterious variants and demographic history of the population jointly determine the degree of its short-term loss while the strength of positive selection determines the rate of trait recovery.

There are substantial differences in the evolutionary dynamics of the Mendelian trait and the polygenic trait under GR. In simulations of a polygenic trait, the consequences of GR on the trait is decided by both the loss of genetic materials as a whole and the distance between the phenotype and its optimum before admixture. Generally speaking, it takes about 100 generations for a polygenic trait to return to its optimum in most cases, which is shorter than that for a Mendelian trait under the same situation. Because polygenic traits have large mutational targets, causal genetic variation that was previously exclusive to the donor population is introduced to the inbred population via GR; this variation quickly replaces native causal genetic variation, which is linked to many deleterious alleles. Thus, the apparently higher efficiency with which the polygenic adaptation is restored comes at the cost of long-term replacement by genetic variation from the donor population. We also showed that the distance of the phenotypic optimum between the donor and the recipient population has appreciable influence on how much genomic replacement is incurred by GR.

Our results demonstrate a marked difference between a long-term small effective population size (Model 1) and a short-term severe bottlenecks (Model 2), with the latter interacting strongly with the dominance of the deleterious mutation load. Our simulations assume that, following GR/admixture, the effective population size of the recipient population immediately recovers to the full size of the ancestral population. Future directions could consider more gradual recoveries in the effective population size, possibly by using evolutionary rescue models such as those discussed by Osmond & Coop (2019). Thus, our models show GR operating at the upper limit of its efficiency, since the aforementioned alternative models would have strictly lower effective population sizes in the short term following admixture.

One caveat of our results is that our simulations do not assume epistasis and, therefore, does not allow for the evolution of Dobzhansky-Muller incompatibilities (DMIs). However, in the presence of DMIs, outbreeding depression may lead to limited genetic replacement or even reduce the absolute fitness after GR.

Although our results suggest that locally adaptive traits, especially those that are Mendelian or moderately polygenic, will be strongly affected by GR in the short term, but the causal variant is generally retained and returns to fixation in the long run. While locally adaptive polygenic traits are less susceptible to shifts due to GR, their underlying genetic architecture is highly susceptible to long-term replacement by donor ancestry.

## Supporting information

zhang2020supp

## Data accessibility

Scripts are available on GitHub at https://github.com/YulinZhang9806/GR_adaptation_scripts.

## Authors contribution

AJS and RN conceptualized the study; YZ and AJS designed the methods; YZ wrote the software; YZ and AJS conducted the analysis; YZ and AJS wrote the manuscript; YZ, AJS, and RN edited the manuscript; AJS and RN supervised the research.

