## Supplementary material for "Evolution of the genetic architecture of local adaptations under genetic rescue is determined by mutational load and polygenicity": zhang2020supp

### Table of Contents:

| | |
|---|---|
| **Simulation settings (Table S1)** | Page 2 |
| **Detailed results-Mendelian trait, hard sweep (Fig S1-S3)** | Page 3-5 |
| **Detailed results-Mendelian trait, soft sweep (Fig S4-S6)** | Page 6-8 |
| **Detailed results-Polygenic trait (Fig S7-S10)** | Page 9-12 |

**Table S1**. Simulation models and parameter settings.

| Adaptive trait type | Selection model | Selection strength | Other settings | Detailed parameters |
|---|---|---|---|---|
| Mendelian | Hard sweep | Selection coefficient ($s$) of the adaptive mutation: $s$=$10^{-2}$, $s$=$10^{-3}$ and $s$=$10^{-4}$ | Dominance coefficient of the deleterious mutations: additive ($h$=0.5), partially recessive ($h$=0.1) and recessive ($h$=0) | <u>*Adaptive mutation:*</u><br>$h$=0.5<br>$s$=$10^{-2}$, $10^{-3}$, $10^{-4}$<br><u>*Deleterious mutations:*</u><br>$h$=0, 0.1, 0.5<br>$s$: picked from a DFE for deleterious mutations[#1] |
| | Soft sweep | | | |
| Polygenic | Stabilizing selection | Strong selection, *Vs*[#2]=3,000 | Optimum before the split of populations, the same as the optimum of outbred population, is 0. Optimum of the inbred population after the split shift by $\delta = 1, 2, 5$ | <u>*Alleles of the polygenic trait:*</u><br>$h$=0.5, $s$=0[#6]<br>*Effects (a):* picked randomly from normal distribution N (0,1)<br>*Phenotype (p)*=$\sum_{i \in SNPs} a_i$[#3]<br>*Fitness*=*calFitness*[#4] * $F(p)$[#5]<br><u>*Deleterious mutations:*</u><br>$h$=0, 0.1, 0.5<br>$s$: picked from a DFE for deleterious mutations[#1] |
| | | Weak selection, *Vs*[#2]=10,000 | | |

[#]1. The DFE used for deleterious mutations are estimated by (Eyre-walker et al. 2006).

[#]2. *Vs* resembles variance in the normal distribution in fitness function.

[#]3. $a_i$ indicates the effect of allele i, phenotype of an individual equals to the sum of the effects of all phenotype-determine loci.

[#]4. *calFitness* here is the fitness contributed by deleterious mutations, calculated multiplicatively with selection coefficients.

[#]5. $F(x)$ is the Probability Distribution Function of normal distribution N (optimum, *Vs*) while $p$ indicates phenotype of an individual.

[#]6. Note that this is for parameter setting in SLiM, it doesn`t mean that the casual SNPs are all neutral.

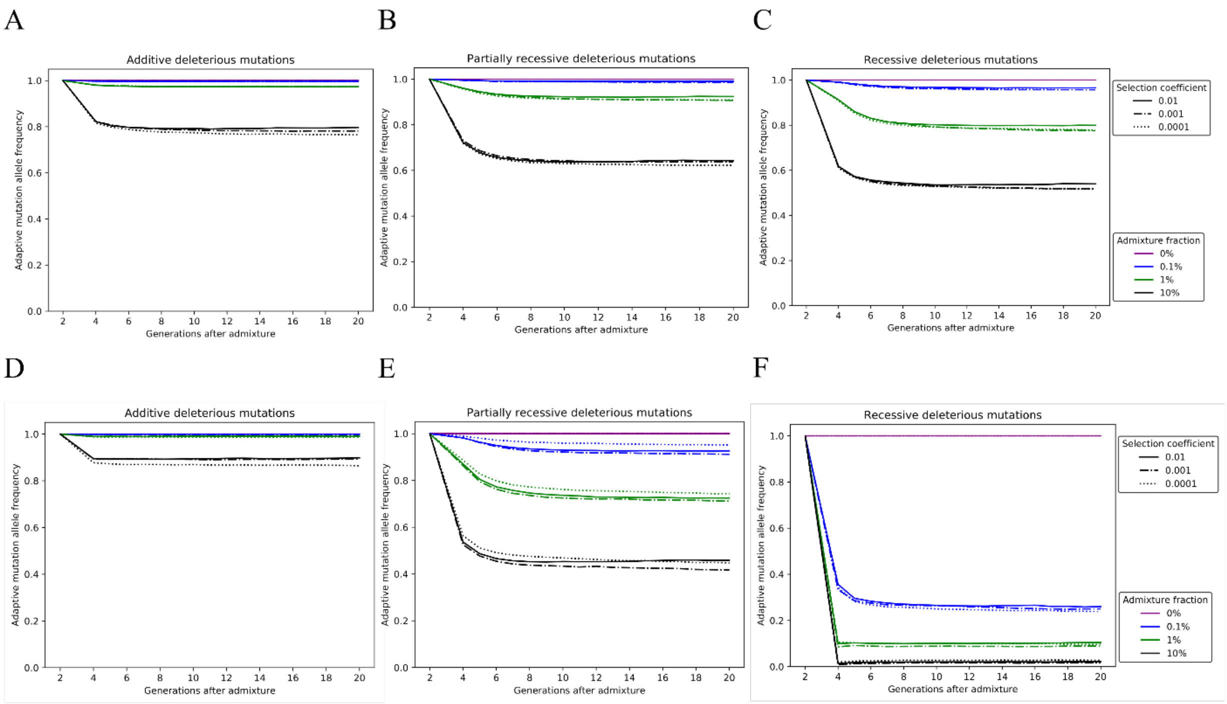


**Figure S1**. Allele frequency changes of 20 generations after admixture for a Mendelian trait under hard sweep selection model and (A-C) demographic Model 1 (D-F) demographic Model 2. The adaptive mutation is additive (dominance coefficient *h*=0.5) while selection coefficient is set differently (shown with different line styles). Deleterious mutations are assumed additive (*h*=0.5) in figure A and D, partially recessive (*h*=0.1) in figure B and E and recessive (*h*=0) in figure C and F.

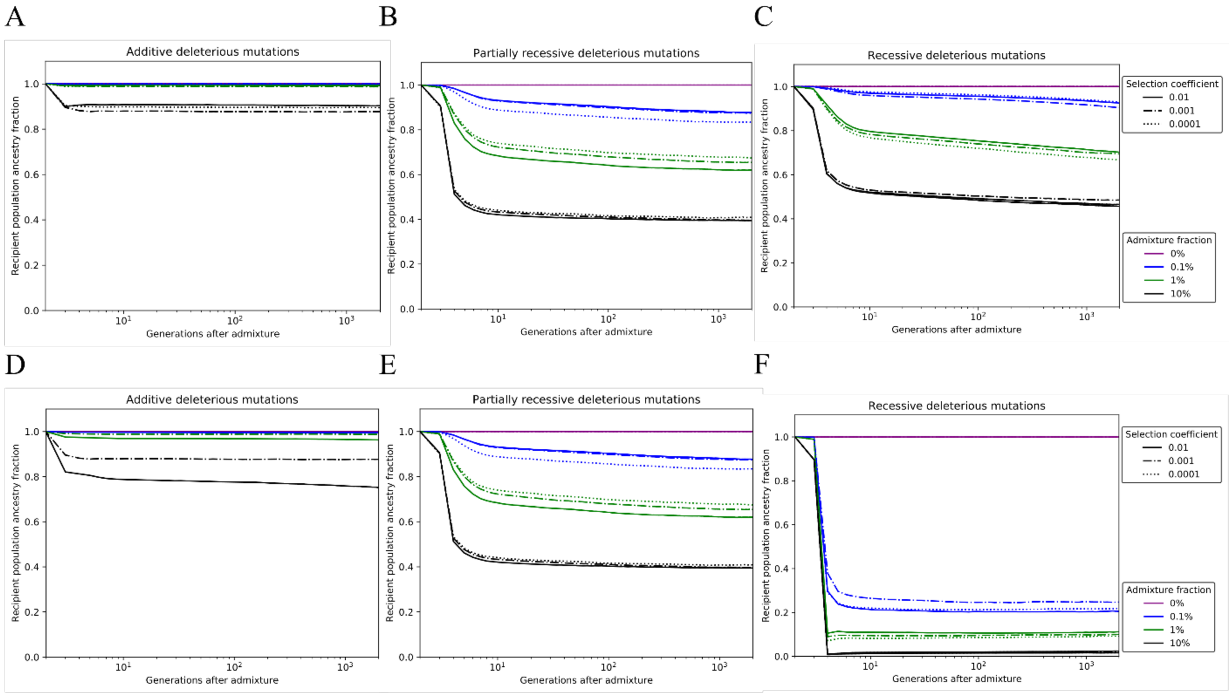


**Figure S2.** Recipient population ancestral genome fraction after GR with a Mendelian adaptive trait, under hard sweep selection model and (A-C) demographic Model 1 and (D-F) demographic Model 2. Each line depicts the ancestral genome proportion of the inbred recipient population. In all scenarios, the adaptive mutation is additive (dominance coefficient *h*=0.5) while selection coefficient is set differently (shown with different line styles). Deleterious mutations are assumed additive (*h*=0.5) in figure A and C, partially recessive (*h*=0.1) in figure B and D and recessive (*h*=0) in figure C and E.

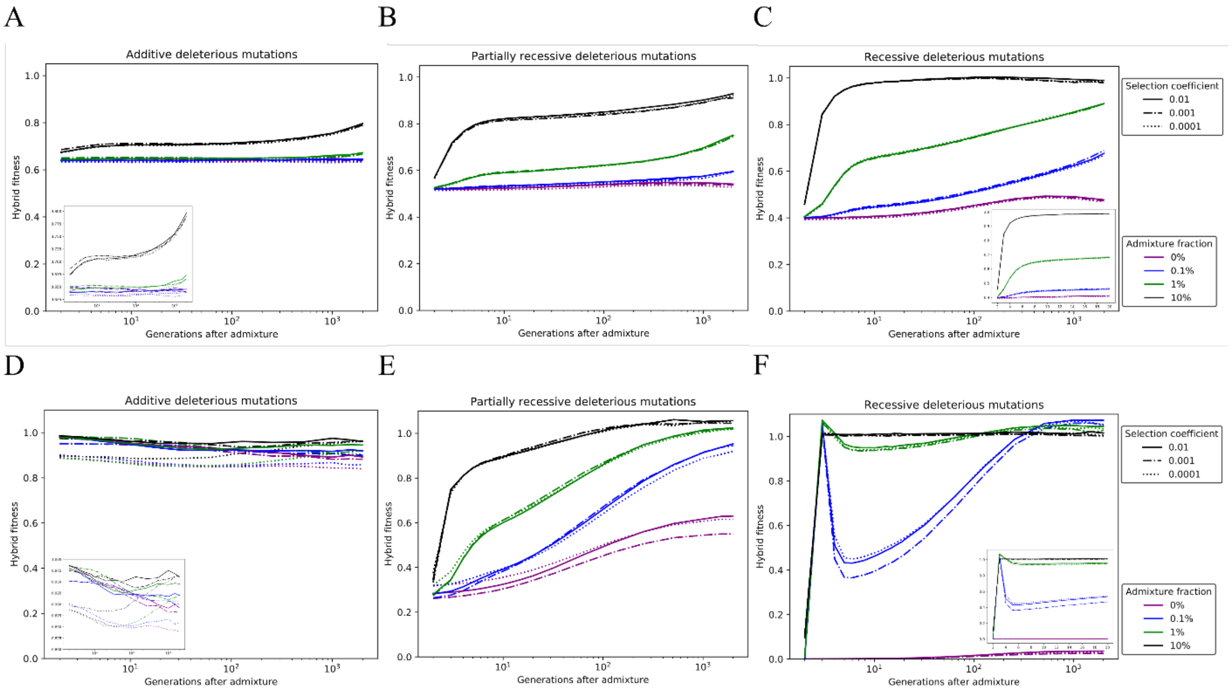


**Figure S3.** Hybrid fitness change of the inbred population after admixture with a Mendelian adaptive trait, under hard sweep selection model and (A-C) demographic Model 1 (D-F) demographic Model 2. The adaptive mutation is additive (dominance coefficient *h*=0.5) while selection coefficient is set differently (shown with different line styles). Deleterious mutations are assumed additive (*h*=0.5) in figure A and D, partially recessive (*h*=0.1) in figure B and E and recessive (*h*=0) in figure C and F.

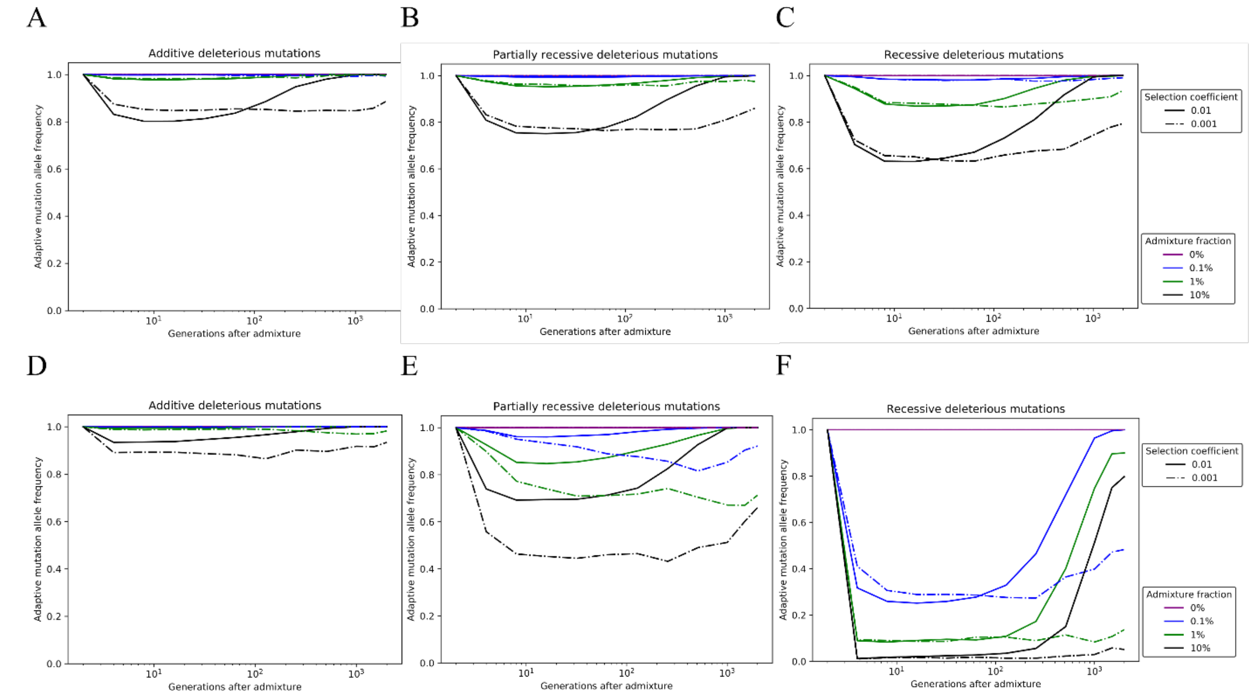


**Figure S4.** Allele frequency changes after admixture for a Mendelian trait under soft sweep selection model and (A-C) demographic Model 1 (D-F) demographic Model 2. The adaptive mutation is additive (dominance coefficient *h*=0.5) while selection coefficient is set differently (shown with different line styles). Deleterious mutations are assumed additive (*h*=0.5) in figure A and D, partially recessive (*h*=0.1) in figure B and E and recessive (*h*=0) in figure C and F.

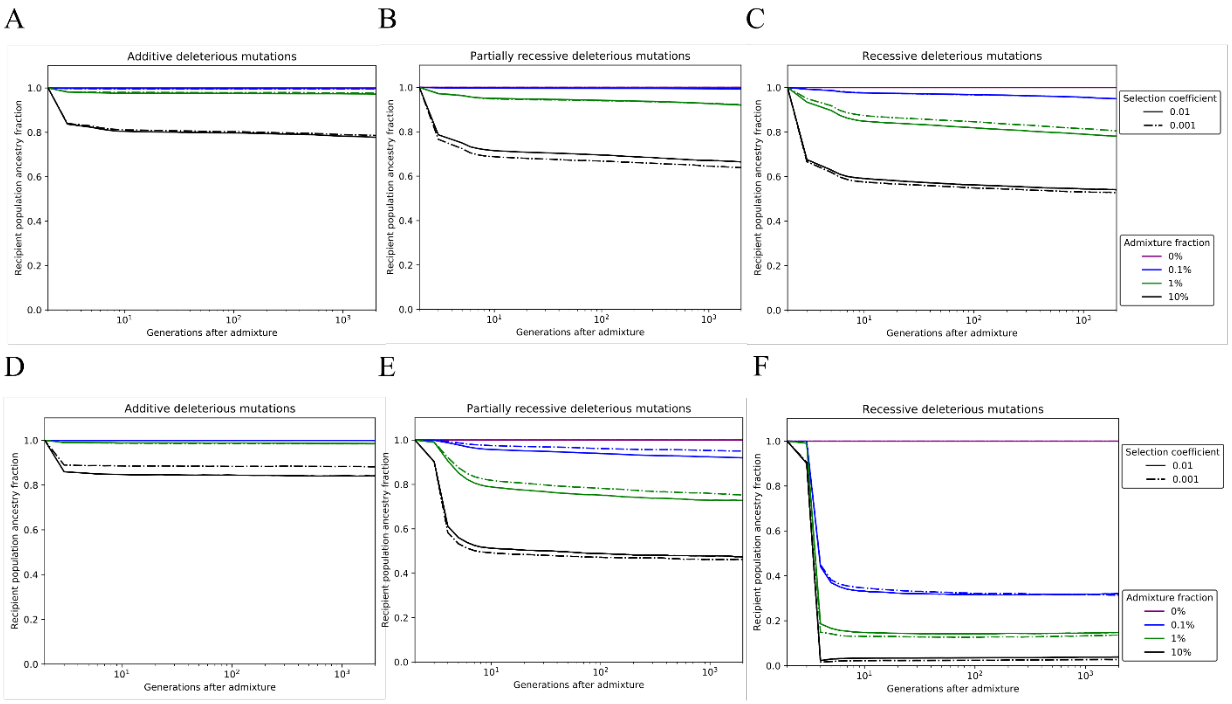


**Figure S5.** Recipient population ancestry fraction after GR with a Mendelian adaptive trait, under soft sweep selection model and (A-C) demographic Model 1 and (D-F) demographic Model 2. Each line depicts the ancestral genome proportion of the inbred recipient population. In all scenarios, the adaptive mutation is additive (dominance coefficient *h*=0.5) while selection coefficient is set differently (shown with different line styles). Deleterious mutations are assumed additive (*h*=0.5) in figure A and C, partially recessive (*h*=0.1) in figure B and D and recessive (*h*=0) in figure C and E.

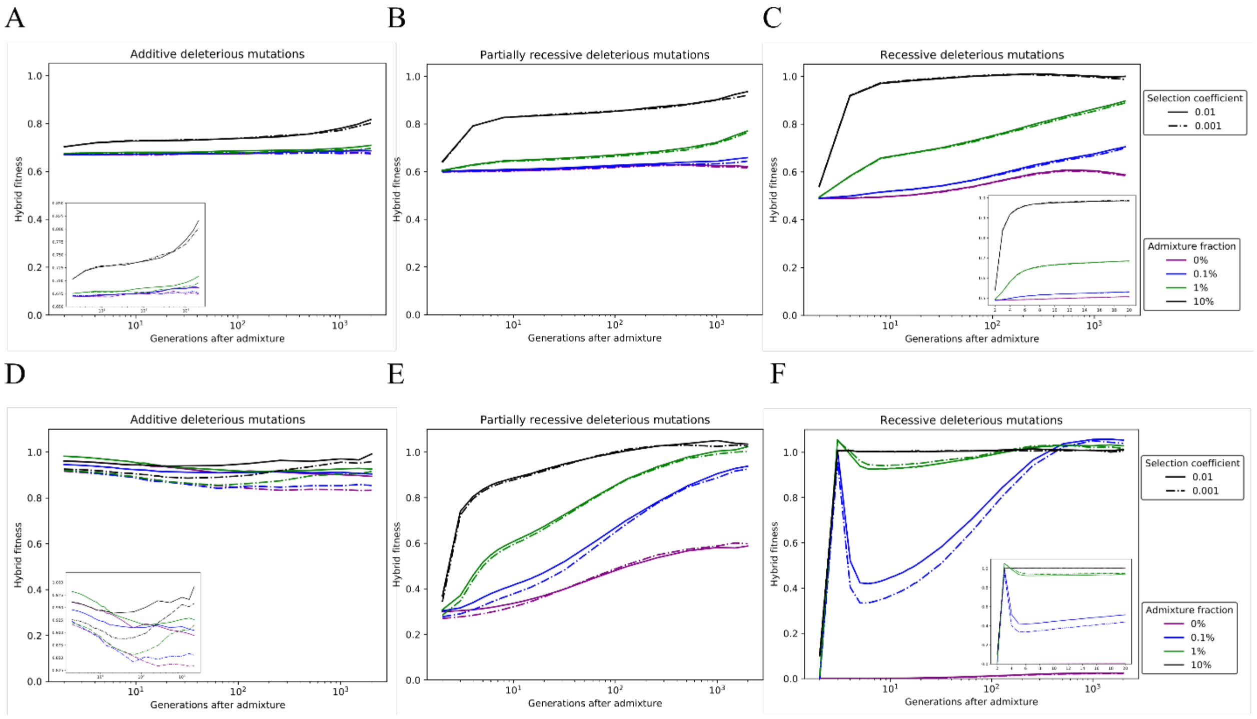


**Figure S6.** Hybrid fitness change of the inbred population after admixture with a Mendelian adaptive trait, under soft sweep selection model and (A-C) demographic Model 1 (D-F) demographic Model 2. The adaptive mutation is additive (dominance coefficient $h$=0.5) while selection coefficient is set differently (shown with different line styles). Deleterious mutations are assumed additive ($h$=0.5) in figure A and D, partially recessive ($h$=0.1) in figure B and E and recessive ($h$=0) in figure C and F.

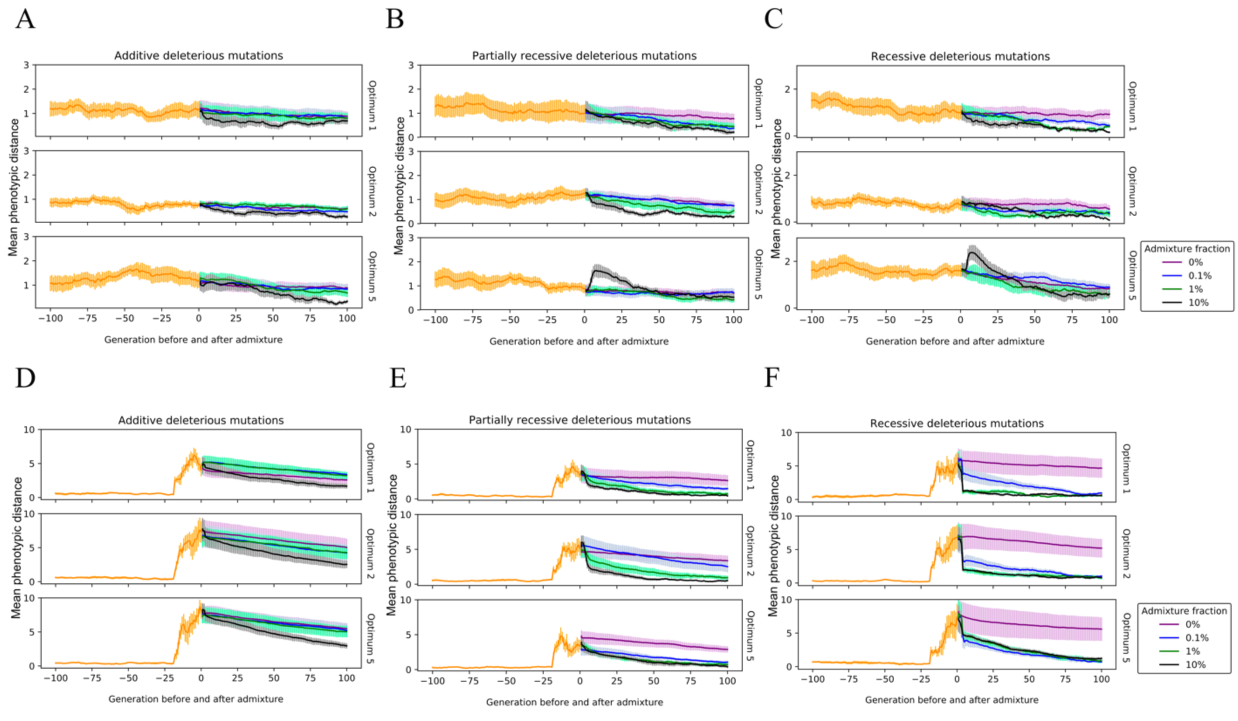


**Figure S7.** Mean phenotype distance from optimum over time. (A-C) Simulations under Model 1, (D-F) simulations under Model 2. Shaded bars signify 95% confidence intervals for the mean phenotypic distance. (Same as Figure 5 in main text, but zoomed into [-100,+100] generations after admixture.)

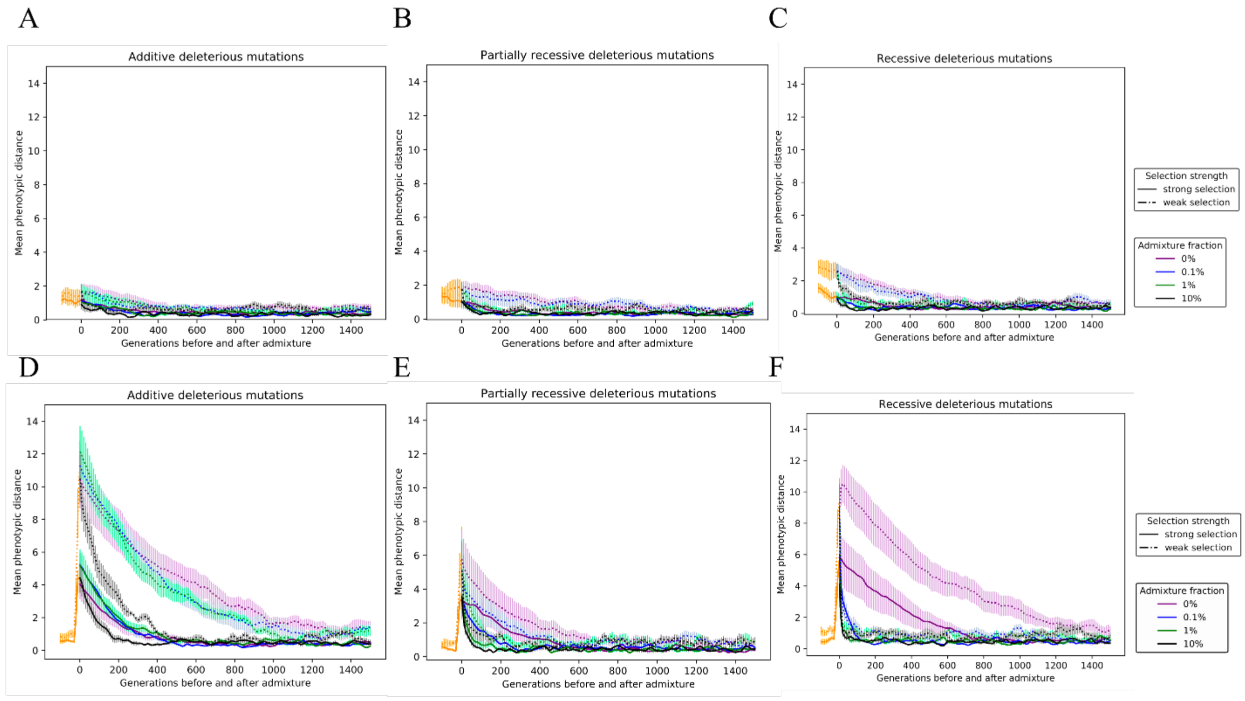

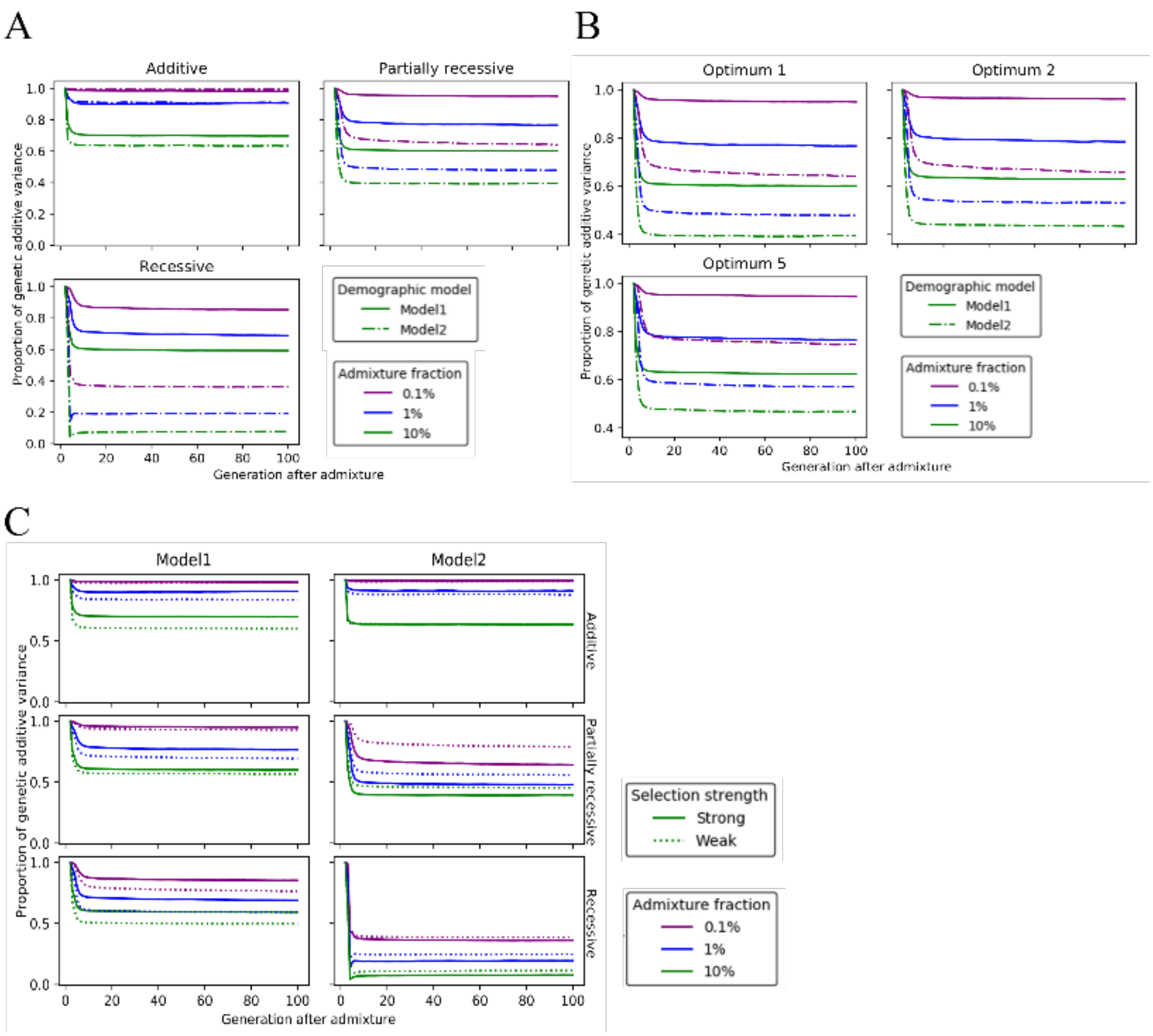


**Figure S8.** Proportion of genetic additive variance of a polygenic adaptation, recording mutations that originate from the recipient population for 100 generations after GR. (A) Scenarios under strong selection, with optimum of 1 for the recipient population and different dominance coefficients for deleterious mutations. (B) Scenarios under strong selection, with partially recessive deleterious mutations and different optimum for the recipient population. (C) Scenarios with optimum equals 1 and different selection strength for the adaptive trait. Each line represents the proportion of genetic additive variance contributed by variants, which compose the trait, from the ancestral genome of inbred population. Here, genetic additive variance=$\sum_{l \in SNPs} 2a_l^2 p_l(1-p_l)$, where $a_l$ represents the effect of SNP $l$, and $p_l$ its frequency.

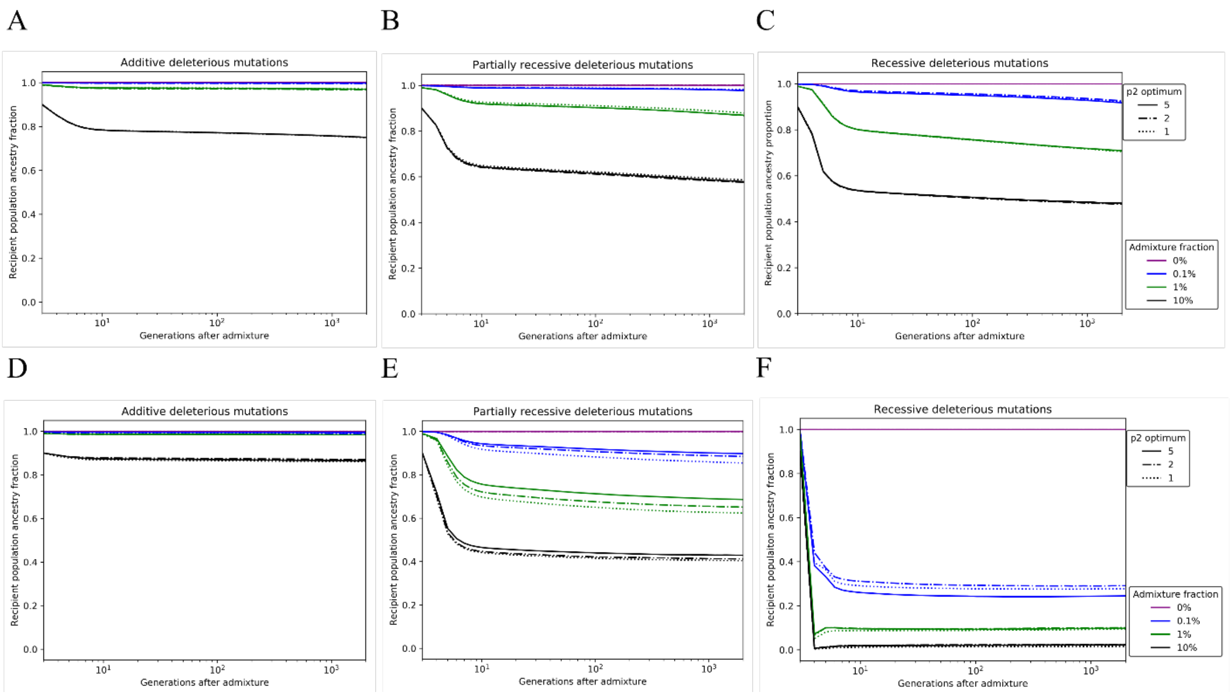


**Figure S9.** Recipient population ancestry fraction after GR with a polygenic adaptive trait, under stabilizing selection model and (A-C) demographic Model 1 and (D-F) demographic Model 2. Each line depicts the ancestral genome proportion of the inbred recipient population. In all scenarios, the adaptive mutation is additive (dominance coefficient *h*=0.5) while phenotypic optimum is set differently (shown with different line styles). Deleterious mutations are assumed additive (*h*=0.5) in figure A and C, partially recessive (*h*=0.1) in figure B and D and recessive (*h*=0) in figure C and E.

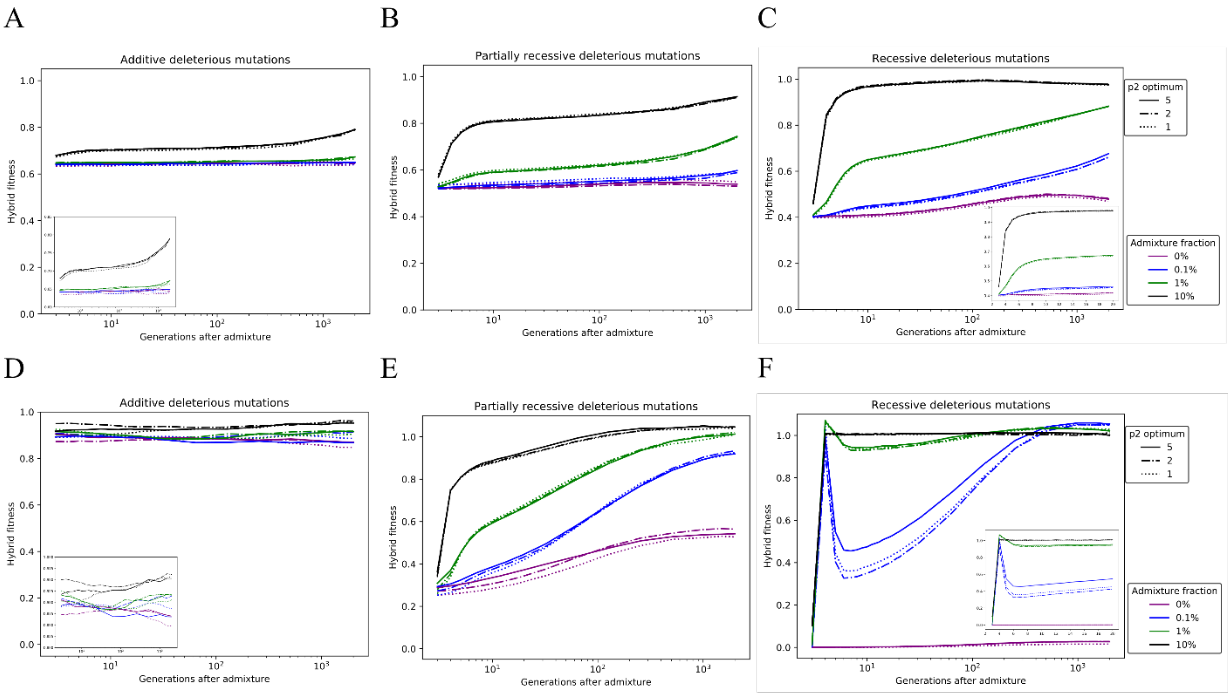


**Figure S10.** Hybrid fitness change of the inbred population after admixture with a polygenic adaptive trait, under stabilizing selection model and (A-C) demographic Model 1 (D-F) demographic Model 2. The adaptive mutation is additive (dominance coefficient $h$=0.5) while phenotypic optimum is set differently (shown with different line styles). Deleterious mutations are assumed additive ($h$=0.5) in figure A and D, partially recessive ($h$=0.1) in figure B and E and recessive ($h$=0) in figure C and F.
